# CRISPRpic: Fast and precise analysis for <u>CRISPR</u>-induced mutations via <u>p</u>refixed <u>i</u>ndex <u>c</u>ounting

**DOI:** 10.1101/416768

**Authors:** Seung Woo Cho, HoJoon Lee, Howard Y. Chang, Hanlee P. Ji

## Abstract

Analysis of CRISPR-induced mutations at targeted loci can be achieved by PCR amplification followed by massively parallel sequencing. We developed a novel algorithm, called CRISPRpic, to analyze sequencing reads from CRISPR experiments via counting exact-matches and pattern-searching. Compared to other methods that are based on sequence alignment, CRISPRpic provides precise mutation calling and ultrafast analysis of sequencing results. The Python script for CRISPRpic is available at https://github.com/compbio/CRISPRpic.

## Background

CRISPR is the most widely used technique for genome editing in research and industry [1]. For successful genome editing in cells and organisms, highly efficient CRISPR-sgRNAs, which differ by cell type or target sequence, are required. Next generation sequencing (NGS) technology allows for convenient and massively parallel measurement of the results of genome editing experiments. Currently, tools exist for analyzing the CRISPR-induced mutation spectrum from sequencing data by aligning sequencing reads to the unmodified reference sequence [2-4]. However, sequence alignment requires multiple calculations for identifying indels with the highest alignment score, and frequently provides false calls depending on the sequence context [5]. We provide a solution to these limitations that offers greater accuracy and speed and is simple to implement.

Herein, we describe an algorithm for fast and precise analysis of <u>CRISPR</u>-induced mutations via <u>p</u>refixed <u>i</u>ndex <u>c</u>ounting (CRISPRpic). Our application has multiple improvements that greatly facilitate detection of variants at CRISPR target sites. We also provide user-friendly customizable input for other mutation analysis applications, i.e. editing using Cpf1. The simplicity of the CRISPRpic algorithm allows for sequencing analysis on a low performance computing environment. Taken together, our method facilitates CRISPR-based experiments, provides greater accessibility to novice researchers who are unfamiliar with the complex nuances of CRISPR-based genome editing, and will increase the analytical throughput of screening for CRISPR-engineered variants across a broad assortment of applications.

## Results

### Overview of CRISPRpic algorithm

We developed a computational tool, CRISPRpic, that counts every possible mutation in a set of sequencing reads without alignment. CRISPRpic is based on three unique properties of gene editing experiments: i) Sequencing reads have fixed ends originating from PCR primer pairs, ii) CRISPR/Cas9 induces a double strand break (DSB) at a predictable position within a target sequence, and iii) Mutations should encompass the DSB site (**Fig. 1a**). These features enable the prediction of the majority of possible mutations and therefore their efficient identification. The default input to the program is the list of amplicon sequences, the guide RNA sequences located within each amplicon, and the type of the endonuclease used, which has a defined breakpoint (CRISPR/Cas9 from different bacterial species: SpCas9 or AsCpf1, etc.). Using these parameters, CRISPRpic has the flexibility to analyze genomic alterations produced by several different enzymes covering a variety of DSB positions. Our pipeline implements the following steps (**Fig. 1a**); i) building a hash table and set of *k*-mer indices, ii) identifying and selecting amplicons with their frequencies, and iii) classifying amplicons by using hash tables or pattern searching of *k*-mers. Most cas9 induced mutations are predictable and stored in hash table for perfect matching. However, some sequencing reads, from some insertions at DSB and unknown variants incorporated by PCR or sequencing error, are not identical to the predictions. In order to classify these reads with unpredictable variants, we designed a pattern searching algorithm using the distance of the *k*-mer indices in the references (m) and sequencing reads (n) (**Fig. 1b**). Basically, the relative difference between m and n indicates insertion, deletion, and so on. Altogether, CRISPRpic is designed to follow a logical decision tree using either exact-matching or pattern-searching, which allows for non-ambiguous mutation calling (**Fig. 2**).

**Fig 1.**
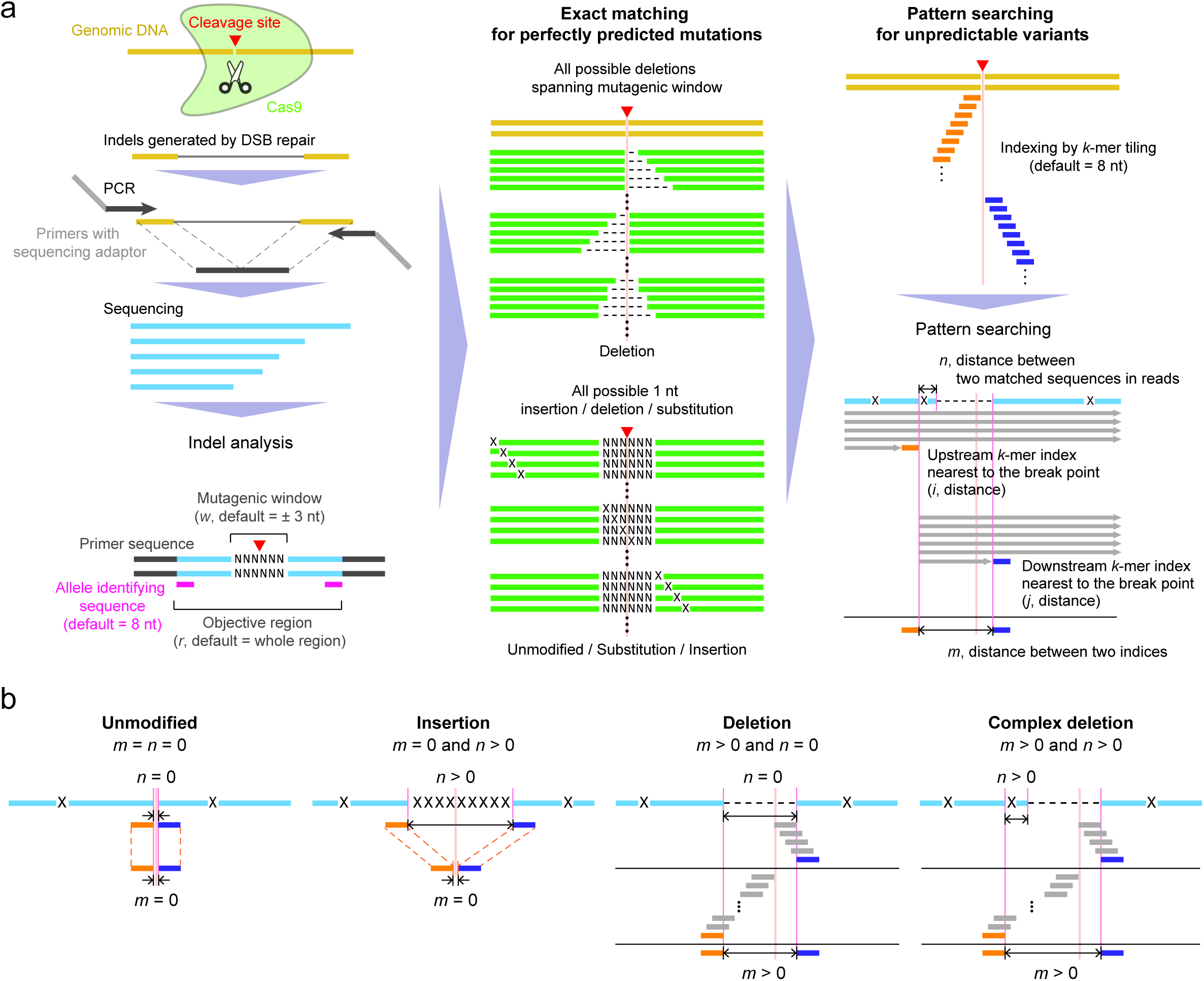
Sequencing analysis of CRISPR-induced mutations using CRISPRpic. **a** Schematics representing the NGS analysis for the CRISPR experiments and CRISPRpic algorithm to analyze them. **b** Examples of mutations calling by pattern-searching algorithm. Reference, sequencing read, or virtual DNA sequence is shown in yellow, blue or green, respectively. Orange and navy indicate the *k*-mer indices.

**Fig 2.**
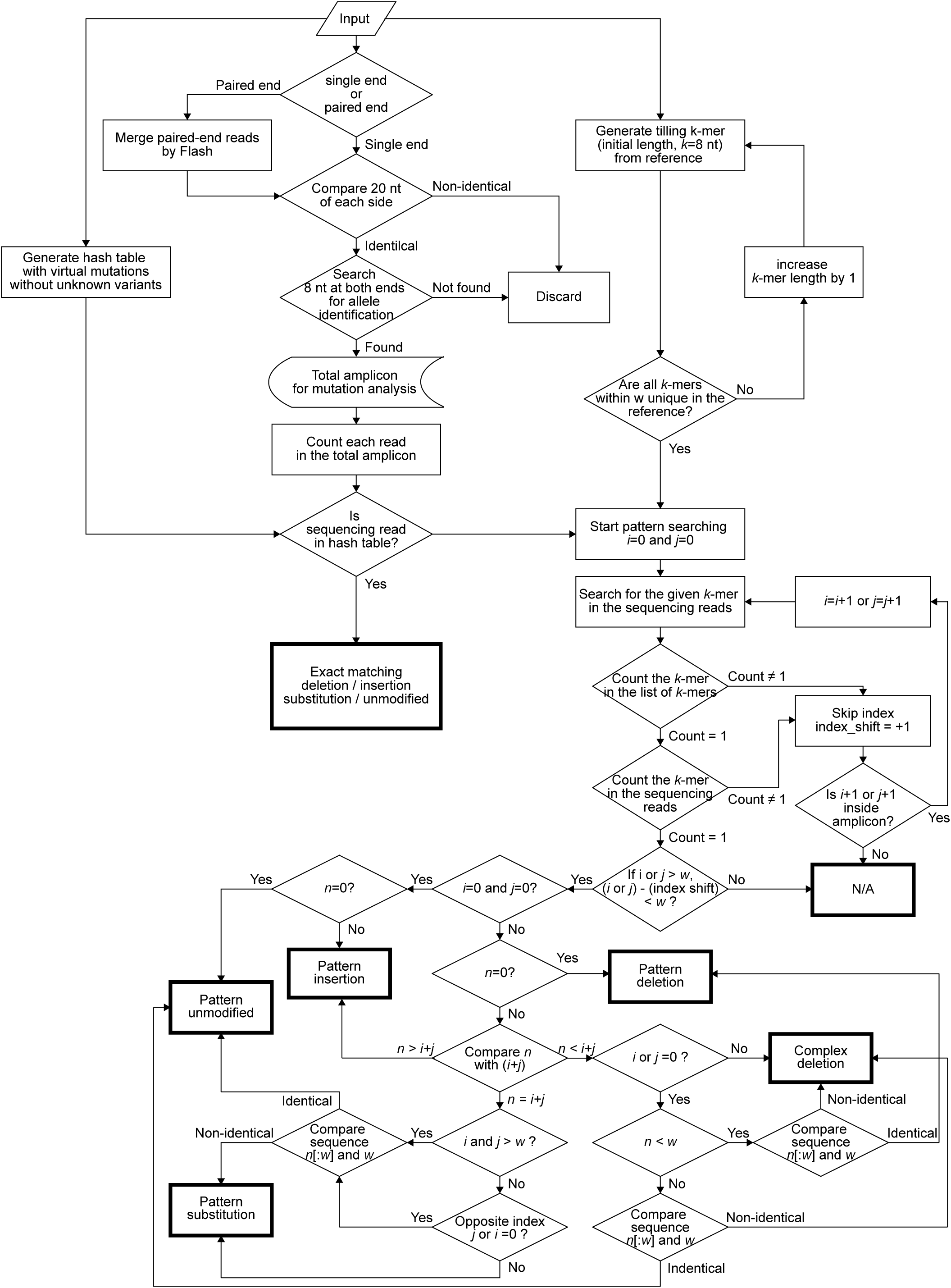
Logical decision tree of CRISPRpic.

### Precise mutation calling by CRISPRpic

We tested CRISPRpic on 20 million reads from the human *PVT1* locus amplified from cells treated CRISPR/SpCas9 [6] (**Fig. 3a**). CRISPRpic successfully classified almost 100% of reads; 94.8% by exact-matching and 5.2% by pattern-matching. Only 0.003% of reads could not be classified appropriately. From this data, we randomly sampled sequencing reads ranging from the 10^3^ to 10^7^ reads. In this analysis, overall mutation frequencies were not significantly different from the total sample size (**Fig. 3b**). As CRISPRpic made all theoretically possible deletions in the hash table, we analyzed the frequencies of each deletion allele. Of the 1,499 deletions that could theoretically occur in this amplicon, we detected 679. (**Fig. 3c**). The frequency of each deletion allele varies with sample size and saturated at 10^7^ reads sample size (**Fig. 3d**).

**Fig 3.**
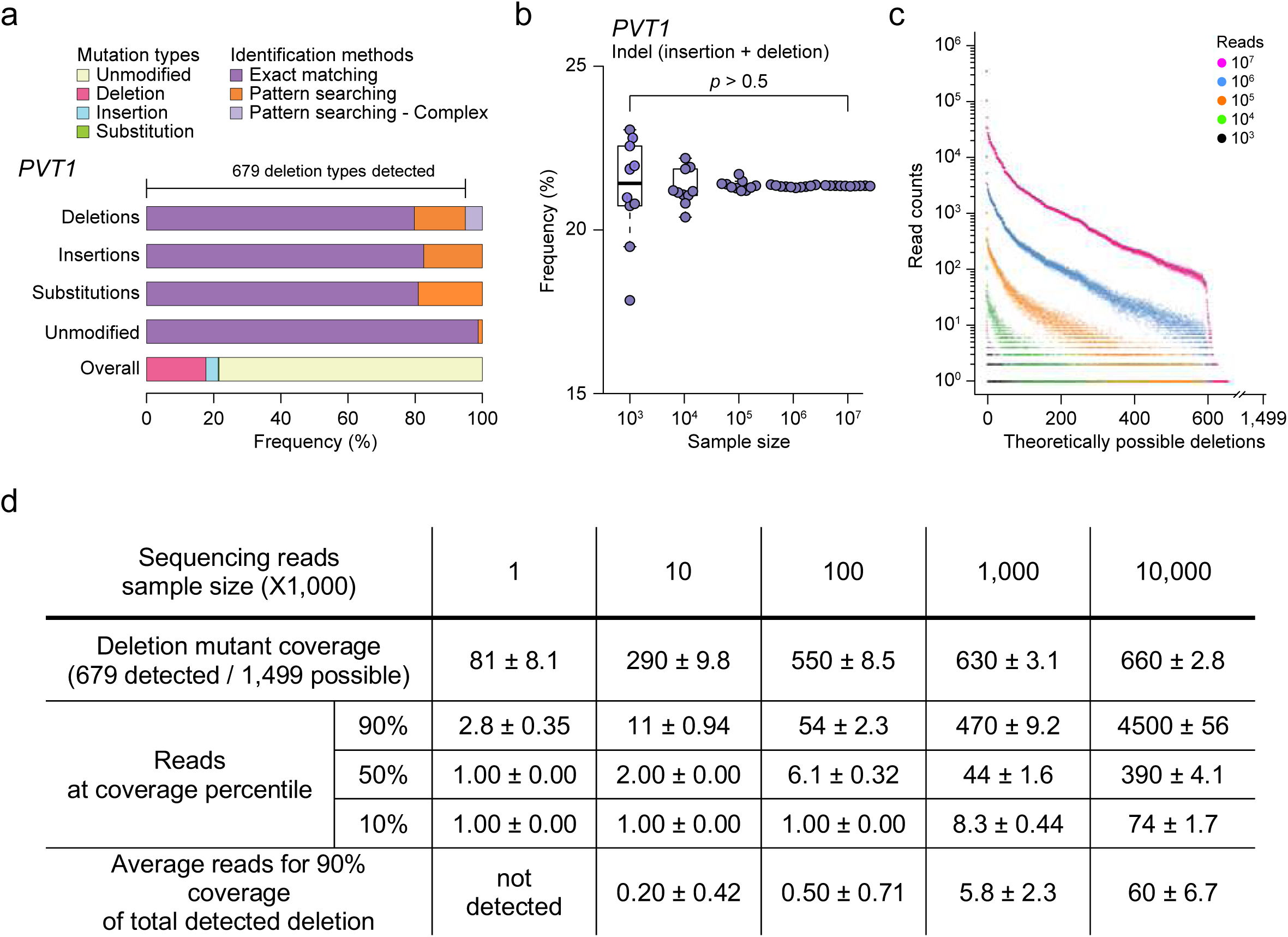
Analysis with different sample size of sequencing reads using CRISPRpic. **a** Mutation analysis of sequencing reads from the *PVT1* locus targeted by SpCas9. **b** Comparison of indel frequencies by sample size of sequencing reads. The *p*-value was calculated by two-sided *t*-test (*n*=10, technical replicates). **c** Distribution of all theoretically possible deletions at the *PVT1* locus by sample size of sequencing reads. All results from repeated sampling are shown together by the sample size. **d** Comparison of i) coverage of all detected deletions, ii) read number at 10th, 50th and 90th percentile of covered deletions, and iii) the read number needed to cover 90% of all detected deletions.

We further analyzed sequencing reads from 10 different loci in the human genome targeted by two different types of CRISPR [7] using three different programs: CRISPRpic, CRISPResso, and Cas-analyzer (**Fig. 4a** and **Additional files 1**). CRISPRpic successfully assigned the vast majority of the sequencing reads (>99.99 % of the total reads at 20 targeted loci) to a single prefixed classification (**Additional files 2**). In this analysis, the three different programs showed significant differences in the indel frequencies for the following loci: *DNMT1*-3, *EMX1*-2, or *HPRT1*-4 of SpCas9, and *CCR5*-1 or *DNMT1*-4 of AsCpf1 (**Fig. 4a**). We found that alignment-based programs showed false mutation calling dependent on the parameters at a particular locus (**Fig. 4b**). We further inspected the reads that were classified differently by the three programs. For the 50 types of sequencing reads classified manually as deletions, CRISPRpic classified them correctly while some of them were not properly classified by the either one of two other programs (**Additional files 3**). Our calls always agreed with majority classification. In addition to deletions, CRISPRpic correctly classified insertions and substitutions. For all reads we examined manually which were classified deletions, insertions or substitutions manually, CRISPRpic classified them correctly, confirming that CRISPRpic is precise and robust to sequencing reads harboring unpredictable artifacts.

**Fig 4.**
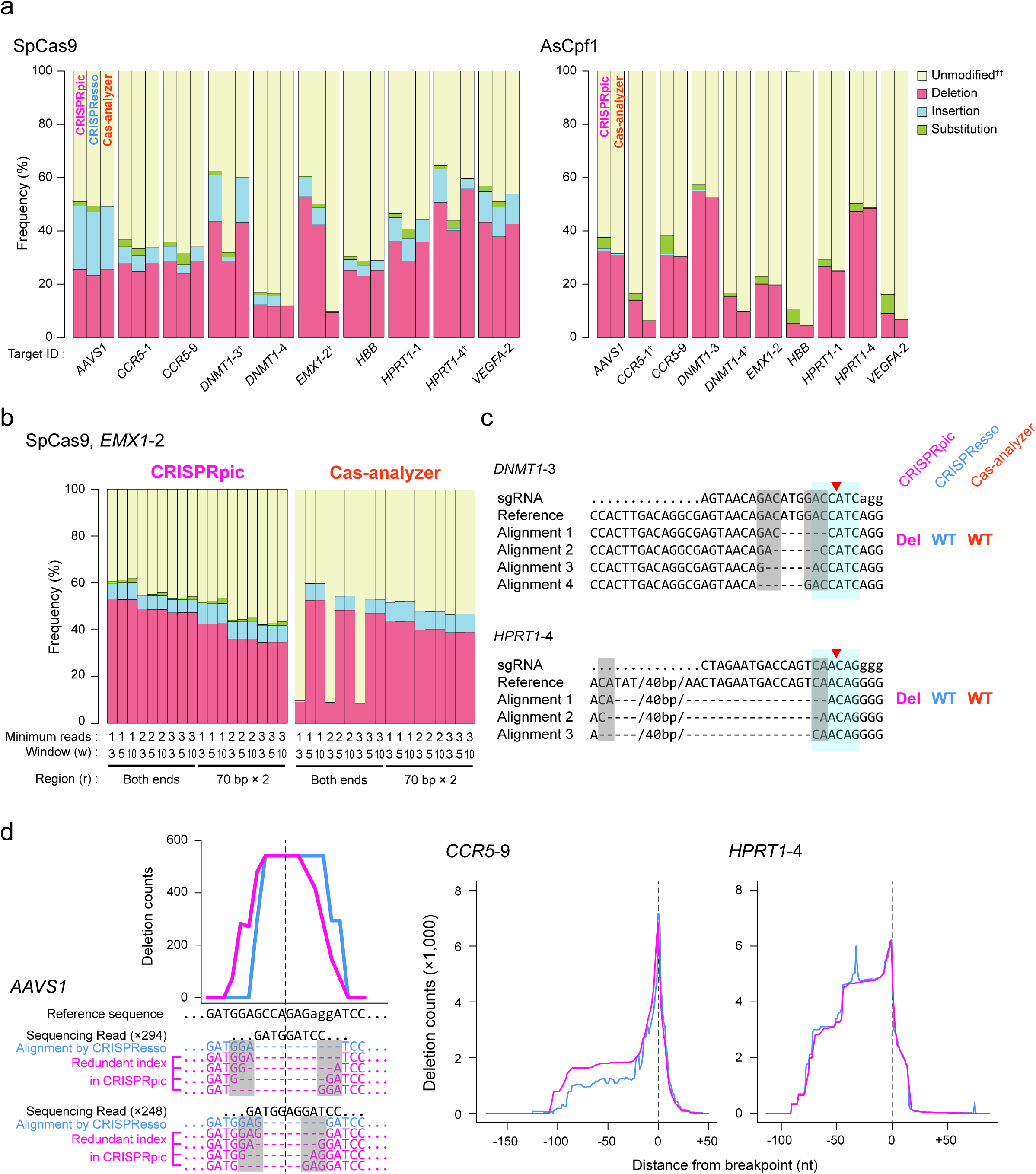
Precise analysis of CRISPR-induced mutations. **a** Mutation frequencies analyzed by CRISPRpic in addition to other programs for 10 different target loci treated by SpCas9 (left) or AsCpf1 (right). **^†^**Sequencing reads were classified manually and more shown in **Additional file 3**. **^††^**Cas-analyzer does not distinguish the substitutions from unmodified allele. **b** Comparison of mutation frequencies by different parameters of CRISPRpic or Cas-analyzer. Colors indicate as shown as **a**. **c** example of classification of deletions harboring micro-homology sequences. Blue or grey box indicates mutagenic window (± 3bp from breakpoint, red triangle) or micro-homology sequences, respectively. **d** Unbiased analysis of deletion pattern by CRISPRpic. Two sequencing reads from the *AAVS1* locus were extracted from actual output files of CRISPRpic (pink) or CRISPResso (blue) as an example (left). Micro-homology sequences are marked in the grey box. Dashed line indicates the breakpoint. Deletion distribution of entire sequencing reads from *CCR5*-9 or *HPRT1*-4 locus (right).

When DSBs are repaired in living cells, micro-homology mediated deletions are more frequent than random ones [8]. The CRISPRpic algorithm can correctively classify mutations harboring micro-homology sequences. By manual examination, we found that alignment-based methods often incorrectly classified the sequences with micro-homology. When micro-homology sequences are present at the border of a mutagenic window, deletions at multiple locations often generate a single sequence. Mutation calling can be ambiguous as their possible multiple alignment and aligning programs generally select one of these possibilities (**Fig. 4c**). Biased alignment of sequence with micro-homology also showed inaccurate distribution of the overall deletion pattern by position (**Fig. 4d**). However, CRISPRpic integrates these possibilities into one deletion entity while preserving multiple deletion position information, successfully explaining the higher frequency of a deletion that can be derived from multiple positions. Therefore, CRISPRpic is robust with micro-homology and precisely presents all possibilities for each deletion in contrast to alignment-based methods which show positional bias in deletion calling.

### Ultrafast analysis of mutations by CRISPRpic

In addition to the precision of mutation calling, CRISPRpic showed drastically decreased analysis time, compared to alignment-based methods (>1,000-fold, **Figure 5a**). Indeed, CRISPRpic is designed to perform simple counting and k-mer searching rather than multiple calculations, thus not requiring a high-performance computing environment. Therefore, CRISPRpic can analyze 20 million reads in only one minute on a personal lap-top computer, whereas CRISPResso could not complete the analysis within 2 days (**Fig 5b**).

**Fig 5.**
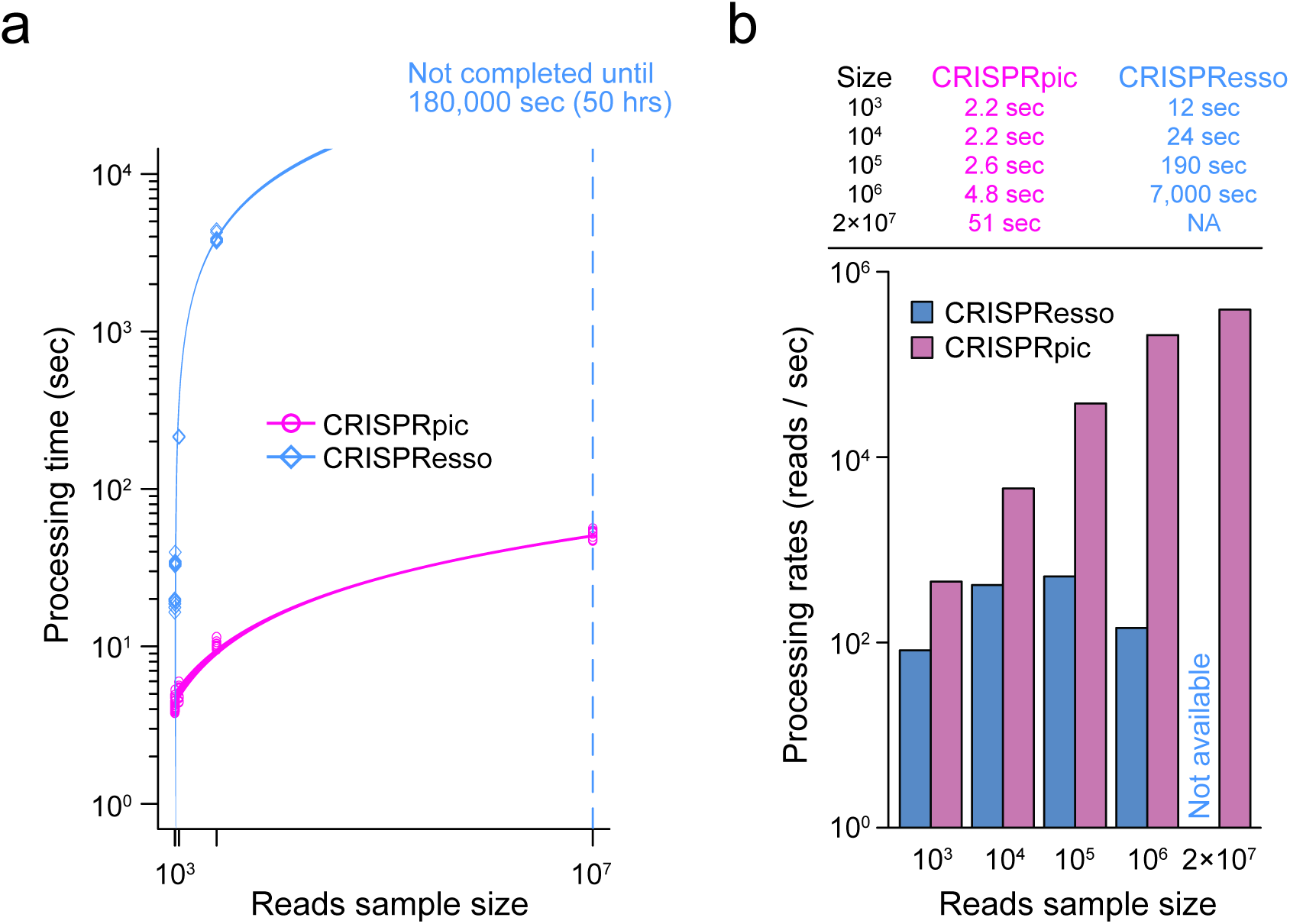
Ultrafast performance of CRISPRpic. **a** Processing time for CRISPRpic and CRISPResso up to 107 sequencing reads from a sample of human PVT1 locus (*n*=10, technical replicates). Lines represent linear regression and 95% confidence intervals. **b** Processing time for mutation analysis on personal lap-top computer using CRISPRpic or CRISPResso.

## Discussion and Conclusions

Induction of mutations at target loci is one of the most common applications of CRISPR, but analysis of the mutation spectrum by sequence alignment has multiple limitations, including incomplete or erroneous calling of variants requirement for a high-performance computing resource. Although large numbers of researchers need to analyze CRISPR-induced mutations, a precise and fast method for analysis has not been developed so far. CRISPRpic is based on exact-matching and logical decisions, which provides precise and ultrafast analysis of mutations in CRISPR experiments.

We note that there are two cases where CRISPRpic is not able to classify (which is classified as NA) and one case where it classifies incorrectly. From our analysis of amplicon sequencing data across 10 loci, we observed only one case as a source of NA; all *k*-mers either upstream or downstream were not found within the amplicons. This can happen a when large deletion left less than k nucleotides at either end or less than (*k* × 2) long nucleotides with some other mutation events. In the analysis of 20 targets, 16 loci did not have any NA and only one locus showed a maximum of 0.04% NA. Sequences labeled as NA were eliminated from initial calculation of indel frequency. CRISPRpic is, in fact, designed to classify an additional case into NA, in which of *i* or *j* index is larger than *w* but [*i* index > *w* and *i* index – *i* shift-count < *w*] or [*j* index > *w* and *j* index – *j* shift-count] is less than *w*. This is the case where it is very challenging to decide clearly whether mutations are within DBS or not. We did not observe this case in the 21 targeted sites we analyzed to develop CRISPRpic.

We found only one type of sequence from the *HPRT1*-4 locus which was incorrectly classified. CRISPRpic called a given set of reads as a complex deletion, but manual examination showed a deletion rather than complex deletion. This incorrect classification occurred because the locus contains an AT-repeat sequence that is longer than the *k*-mers in the upstream sequence. In this case, CRISPRpic skipped several non-unique *k*-mers due to AT repeats resulting in *i* is not zero followed by classification into a complex deletion. For cases like this, in which the target locus contains repetitive sequences, we recommend that users provide a longer *k*-mer length as an input parameter.

CRISPRpic is useful for high-throughput screening of efficient sgRNAs for CRISPR, as well as for other applications such as mutagenesis-based functional studies of proteins [9] and regulatory elements in a DNA-centric manner [10]. The advent of single cell analysis of CRISPR-mediated perturbations [11-13] further necessitates the need for highly efficient and scalable means of analyzing gene edits. From the analysis conducted in this study, we recommend sequencing 10^3^ to 10^4^ sequencing reads per target locus to accurately evaluate the efficiency of CRISPR. For allele-based quantitative analysis, we recommend sequencing 10^6^ to 10^7^ total reads depending on the mutation frequency (**Fig. 3**).

Recently, it was reported that CRISR can induce large deletions over several kilobases [14]. When surveying long range mutations, longer PCR combined with sequencing errors can cause false alignment-based calls followed by incorrect mutation calling. CRISPRpic is exceptional at distinguishing variants not induced by CRISPR, which will make CRISPRpic a standard method for analysis of CRISPR-induced mutations for any type of amplicons.

CRISPRpic is precise and robust with sequence context, thus will provide a clear picture of Cas9 induced mutations in CRISPR-based experiments. CRISPRpic is also ultrafast, so enables researchers to analyze the large size of sequences data from CRISPR-based genome editing, which is not feasible by any other tools.

## Methods

### Detailed algorithm of CRISPRpic

Step1: Building hash tables for exact matching

CRISPRpic builds a hash table representing all of the sequences with all possible mutations encompassing the target site of DBS in the reference sequence, which is the expected amplicon. This target site, or the CRISPR mutagenic site, is determined by the window size (*w*, default = 3) defined by users. Therefore, the mutagenic site encompasses the breakpoint with 3 nucleotides on both sides, leading to a total 6 nucleotides with the breakpoint in the middle (**Fig. 1a**). We consider only mutations within this mutagenic site as CRISPR-induced mutations. All other mutations (which tend to be PCR artifacts or sequencing errors) are not considered as mutations induced by Cas9.

First, all possible deletions encompassing the mutagenic window and with different sizes are generated (**Fig. 1a**). Some deletions at different positions can be identical when short, common sequence motifs (i.e. micro-homology regions) are present in the reference sequence. In this case, we treat them as single deletion event with multiple alignment positions (**Fig. 4d**). Second, the hash table includes single nucleotide substitutions, deletions, and insertions at all positions of the reference sequence (**Fig. 1a**). Only substitutions, deletions, and insertions within the defined CRISPR mutagenic window are classified as mutations while all other outside changes are classified as “unmodified”.

Step 2: Building two sets of k-mer indices for pattern searching

Some portion of amplicons cannot be matched to one of sequences in the hash table by exact matching when they have unknown variants not induced by endonuclease. To classify them, we employ pattern searching using a *k*-mer “index” (default length of *k* is 8). The program generates a set of *k*-mer indices from the middle breakpoint and tiles upstream in 1 nucleotide increments (**Fig. 1a**). Therefore, the order of indices represents the distance from the breakpoint. For example, the first index among the upstream set is located at the right at the breakpoint while second index locates the one nucleotide upstream of breakpoint. Next, we examine the unique representation of each *k*-mer index among all *k*-mer indices. If a *k*-mer appears more than once, we skip this *k*-mer index for pattern searching given there is a possibility of a false positive.

If the outside index is originated from inside the window by skipping, then classification is challenging. We keep all indices within the mutagenic window to facilitate the analysis. Thus, we increase the length of the *k*-mer by one nucleotide until all indices within the mutagenic window are unique. For instance, all three *k*-mer indices must be unique among all indices in upstream set when the size of mutagenic window is 3 We build the downstream set of *k*-mer indices in the same way.

Step 3: Identifying and selecting amplicons

The input data for CRISPRpic are single reads in FASTQ formats that originate from PCR amplicons. The paired-end reads can be converted to single-end reads by a program called FLASH. We process only those reads per the following criteria: they contain either one of two adaptor sequences; both the first and last eight nucleotides of the reference sequence are present after removing adaptor sequences. Based on this processing, we determine the individual amplicon reads with their fractions among the total data set.

Step 4: Classifying amplicon read sequences

First, we examine if all distinctive amplicons are identical to one of the virtual mutant sequences and the reference sequence in the hash table. If they are matched, they will be characterized by the classification in the hash table. Second, all reads not identical to one of sequences in hash table will be classified by pattern searching (**Fig. 1a**). The classification is determined by the following 5 variables with the given mutagenic window size:

1. *i* index, ordinal number of upstream *k*-mer index, which was first found in the amplicon
2. *i* shift-count, number of upstream *k*-mer skipped
3. *j* index, ordinal number of downstream *k*-mer index, which was first found in the amplicon
4. *j* shift-count, number of downstream *k*-mer skipped
5. *n*, the length of remaining sequence between two *k*-mer in the amplicon

The ordinal number of each index indicates the distance from breakpoint. For example, the ordinal number of first index is zero, which represents the right breakpoint on the reference. CRISPRpic initiates a search from the upstream *k*-mer in order until a *k*-mer is found in the amplicon. As mentioned above, we skip the non-unique *k*-mers in the amplicon. The number of skipped *k*-mer are counted as *i*-shift count. CRISPRpic repeats the same procedure for downstream *k*-mer. When no proper *k*-mer is found either upstream or downstream index set on amplicon, we designate this read as “NA”. After finding upstream and downstream *k*-mer in amplicon, CRISPRpic examines the sequences between two identified *k*-mer in amplicons.

After identifying the five parameters above, we eliminate those amplicon sequences when they have [*i* index > *w* and *i* index – *i* shift-count < *w*] or [*j* index > *w* and *j* index – *j* shift-count]. This step is take because for this specific case, mutation identification is a challenge. Otherwise, we classify reads by rules depicted in logical flow chart (see below) using the following three numbers: *i* (= *i* index – *i* shift-count), *j* (= *j* index – *j* shift-count) and *n*.

Step 5: Variant classification

We describe how these parameters are used for identifying a variant using the parameters as described (**Fig. 1b and Fig. 2**). Each case is considered as follows:

i. Reads are classified as reference “wildtype” and not CRISPR-modified When *i, j* and *n* equal zero. Described simply, the first *k*-mer in both upstream and downstream were found and there is nothing between these two *k*-mer in amplicon. This also means 16 nucleotides centered at the breakpoint (i.e. the targeted mutagenic site) are identical to the reference sequence. These sequences are not matched to one of all possible mutated sequences because of unknown variants somewhere outside of mutagenic window of amplicon.
ii. Reads are classified as an insertion When *i* and *j* equal zero, but *n* is larger than zero. This happens when some sequences are inserted at the breakpoint. In this case, first upstream and downstream *k*-mers will be found, but there will be remaining sequence between two indices (*n* > 0) in amplicon due to insertion.
iii. Read are classified as a deletion When *n* equals zero, but *i* or *j* is not equal to zero. This indicates some sequences are deleted at the breakpoint. For example, 2nd upstream *k*-mer and 4th downstream *k*-mer will be found when 1 and 3 nucleotides are deleted from upstream and downstream of breakpoint respectively. However, there will be no remaining sequence between two *k*-mer on amplicon. Finally, there are complicated cases that *m* > 0 and *n* >0, where *m* = *i* + *j*. The classification is determined by the rules in the logic flow chart.
iv. For complex variants that do not fall in the aforementioned categories on the first analysis pass We compare the expected distance (*m*) of two found *k*-mer in the reference sequence with the length of remaining sequence (n). There are three possibility between *m* and *n*; i) *n* > *m*, ii) *n* = *m*, and iii) *n* < *m*.

First, amplicons are classified as insertion when *n* is larger than *m*. This happens when insertion happens with some other events. Second, amplicons are classified as either substitution or unmodified when *n* equals *m*. In this case, we examine if the remaining sequence is identical to the sequence with the mutagenic site. If they are identical, amplicons are classified as unmodified. Otherwise, they are classified as substitution. Third, amplicons are classified as either complex deletion or unmodified when *n* is smaller than *m*. In this case, we examine the location of deletion relative to breakpoint. If deletion occurs outside of the mutagenic site, they are classified as complex deletion. Otherwise, they are classified as unmodified.

### Calculating the frequency of deletions at all position in the amplicon

Multiple deletion events at different position of the reference sequence results in same sequence when there are “micro-homology” sequences at the junction of deletion (**Fig. 4c**). The micro-homology refers to a short sequence motif that appears repeatedly, thus providing multiple alignments to the reference sequence. We use the count of this deletion sequence for each position by dividing by total number of multiple alignments that contain the position. For instance, we observed a deletion sequence 100 times that derived from five different alignments of deletions. All positions deleted in all five alignments are counted 100 times as well. However, the deletion at the most upstream can occur in only one alignment, so it was counted only 20 times (= 100/5).

These deletions as counted though micro-homology motifs tend to be more prevalent than a random deletion that occurs because of sequencing artifact. We use the presence or absence of the micro-homology motif to determine the frequency of out-of-frame deletions. As has been previously reported, this analysis is based on the sequence interval that lies between two micro-homology motifs[8].

### Information of computing environment

All analysis was performed under following specification except **Fig. 5b:**

2.5 GHz AMD Opteron 6380, 512 GB 1600Mhz DDR3, Linux 4.4.0-122 generic #146-Ubuntu SMP

Low performance computing environment for **Fig. 5b:**

2.3GHz Intel Core i5, 8 GB 2133 MHz LPDDR3, macOS High Sierra 10.13.4.

### Software version used in this study

Python 2.7

Flash 1.2.11

CRISPResso, 1.0.8

(with -w 3 -a [amplicon sequence] -g [target sequence-without PAM-])

Cas-Analyzer, web version (http://www.rgenome.net/cas-analyzer/) (with [checked in the ‘or use both ends’], [1 for minimum frequency], [3 for WT marker] for all analysis except **Fig. 4b**)

### Manual inspection for comparison between programs

When manually inspected, the inspector could not determine clearly a specific variant call in many cases. To eliminate this ambiguity we developed the following rules for assignment: i) Mutations are only classified as unmodified, insertion, substitution, or deletion; and ii) If the sequencing read is assigned in multiple categories, we prioritize insertions or deletions above wildtype or substitution. For the CRISPResso that does not provide the classification, we assigned the classification to reads using logical values and numbers from the following columns: NHEJ, UNMODIFIED, n_deleted, n_inserted, and n_mutated. As a test, this manual inspection was done in a blinded fashion.

## List of abbreviations

NGS: Next Generation Sequencing
CRISPR: Clustered Regularly Interspaced Short Palindromic Repeats
DSB: double strand break
sgRNA: single guide RNA
SpCas9: Streptococcus pyogenes Cas9
AsCpf1: Acidaminococcus sp. Cpf1

## Declarations

The authors declare following competing interests: The authors are preparing a file for an intellectual property based on this study.

## Availability of data and materials

CRISPRpic is implemented in Python 2.7 or 3.4 and the source code are available at Github (https://github.com/compbio/CRISPRpic).

All raw sequencing reads used in this study are available through NCBI Sequencing Read Archive with accession number SRP070794[7] and Gene Expression Omnibus with accession number GSM2572600[6].

## Authors’ contributions

All authors conceived and designed the study. S.W.C., HJ.L. and H.P.J. designed the algorithm and analyzed the data. HJ.L. wrote the Python script. All authors wrote the manuscript together. H.Y.C and H.P.J supervised the project.

## Acknowledgements

We thank to Stephanie Greer for her help in testing and depositing CRISPRpic to Github. We also thank to Ahmed Shuaibi, Jiamin Chen, Jing He, Soah Lee, Sue Grimes, and Grace Chen for their helps in testing CRISPRpic. We appreciate Ava C. Carter helped in preparing this manuscript.

## Additional files

Additional files 1. Target locus and amplicon sequences.

Additional files 2. Raw data of mutation analysis using CRISPRpic for 20 different targets.

Additional files 3. Manual analysis of most abundant 10 sequencing reads classified differently by other programs.

## REFERENCES

1. Kim H, Kim JS: A guide to genome engineering with programmable nucleases. Nat Rev Genet 2014, 15:321–334.

2. Pinello L, Canver MC, Hoban MD, Orkin SH, Kohn DB, Bauer DE, Yuan GC: Analyzing CRISPR genome-editing experiments with CRISPResso. Nat Biotechnol 2016, 34:695–697.

3. Park J, Lim K, Kim JS, Bae S: Cas-analyzer: an online tool for assessing genome editing results using NGS data. Bioinformatics 2017, 33:286–288.

4. Guell M, Yang L, Church GM: Genome editing assessment using CRISPR Genome Analyzer (CRISPR-GA). Bioinformatics 2014, 30:2968–2970.

5. Lunter G, Rocco A, Mimouni N, Heger A, Caldeira A, Hein J: Uncertainty in homology inferences: assessing and improving genomic sequence alignment. Genome Res 2008, 18:298–309.

6. Cho SW, Xu J, Sun R, Mumbach MR, Carter AC, Chen YG, Yost KE, Kim J, He J, Nevins SA, et al: Promoter of lncRNA Gene PVT1 Is a Tumor-Suppressor DNA Boundary Element. Cell 2018, 173:1398–1412 e1322.

7. Kim D, Kim J, Hur JK, Been KW, Yoon SH, Kim JS: Genome-wide analysis reveals specificities of Cpf1 endonucleases in human cells. Nat Biotechnol 2016, 34:863–868.

8. Bae S, Kweon J, Kim HS, Kim JS: Microhomology-based choice of Cas9 nuclease target sites. Nat Methods 2014, 11:705–706.

9. Findlay GM, Boyle EA, Hause RJ, Klein JC, Shendure J: Saturation editing of genomic regions by multiplex homology-directed repair. Nature 2014, 513:120–123.

10. Vierstra J, Reik A, Chang KH, Stehling-Sun S, Zhou Y, Hinkley SJ, Paschon DE, Zhang L, Psatha N, Bendana YR, et al: Functional footprinting of regulatory DNA. Nat Methods 2015, 12:927–930.

11. Adamson B, Norman TM, Jost M, Cho MY, Nunez JK, Chen Y, Villalta JE, Gilbert LA, Horlbeck MA, Hein MY, et al: A Multiplexed Single-Cell CRISPR Screening Platform Enables Systematic Dissection of the Unfolded Protein Response. Cell 2016, 167:1867–1882 e1821.

12. Dixit A, Parnas O, Li B, Chen J, Fulco CP, Jerby-Arnon L, Marjanovic ND, Dionne D, Burks T, Raychowdhury R, et al: Perturb-Seq: Dissecting Molecular Circuits with Scalable Single-Cell RNA Profiling of Pooled Genetic Screens. Cell 2016, 167:1853–1866 e1817.

13. Kalhor R, Mali P, Church GM: Rapidly evolving homing CRISPR barcodes. Nat Methods 2017, 14:195–200.

14. Kosicki M, Tomberg K, Bradley A: Repair of double-strand breaks induced by CRISPR-Cas9 leads to large deletions and complex rearrangements. Nat Biotechnol 2018.

